# Recovering rearranged cancer chromosomes from karyotype graphs

**DOI:** 10.1101/831057

**Authors:** Sergey Aganezov, Ilya Zban, Vitaly Aksenov, Nikita Alexeev, Michael C. Schatz

## Abstract

Many cancer genomes are extensively rearranged with highly aberrant chromosomal karyotypes. Structural and copy number variations in cancer genomes can be determined via abnormal mapping of sequenced reads to the reference genome. Recently it became possible to reconcile both of these types of large-scale variations into a karyotype graph representation of the rearranged cancer genomes. Such a representation, however, does not directly describe the linear and/or circular structure of the underlying rearranged cancer chromosomes, thus limiting possible analysis of cancer genomes somatic evolutionary process as well as functional genomic changes brought by the large-scale genome rearrangements.

Here we address the aforementioned limitation by introducing a novel methodological framework for recovering rearranged cancer chromosomes from karyotype graphs. For a cancer karyotype graph we formulate an Eulerian Decomposition Problem (EDP) of finding a collection of linear and/or circular rearranged cancer chromosomes that are determined by the graph. We derive and prove computational complexities for several variations of the EDP. We then demonstrate that Eulerian decomposition of the cancer karyotype graphs is not always unique and present the Consistent Contig Covering Problem (CCCP) of recovering unambiguous cancer contigs from the cancer karyotype graph, and describe a novel algorithm `CCR` capable of solving CCCP in polynomial time.

We apply `CCR` on a prostate cancer dataset and demonstrate that it is capable of consistently recovering large cancer contigs even when underlying cancer genomes are highly rearranged. `CCR` can recover rearranged cancer contigs from karyotype graphs thereby addressing existing limitation in inferring chromosomal structures of rearranged cancer genomes and advancing our understanding of both patient/cancer-specific as well as the overall genetic instability in cancer.

## Background

Cancer is a deadly disease propagated by accumulation of genetic mutations that range across scales from single nucleotide polymorphisms to large-scale genome rearrangements. Advances in whole-genome sequencing of tumor samples have enabled the detection of somatic mutations using specialized algorithms to identify different classes of mutations from short, long, and linked DNA sequence reads available by current technologies [14, 15, 12, 21, 13, 26, 22, 16, 5, 8].

In this study we focus on large-scale genome rearrangements that accumulate throughout the somatic evolutionary process to alter the order, orientation, and quantity of genomic sequences, and ultimate producing a collection of highly rearranged chromosomes that constitute the cancer genome. Large-scale alterations of the chromosomal structure in cancer genomes have been previously identified as risk factors [25], survival prognosis predictors [19, 10], and possible therapeutic targets [23, 18], thus underscoring the importance of this type of mutation and prompting the development of novel methods to decipher the rearranged structures of cancer genomes.

Previously proposed algorithms enabled the inference of *copy number aberrations* (CNA) (e.g., deletion, amplification) of genomic segments [8, 27, 7] and *novel adjacencies* (NA) between genomic loci that are distant in the reference but are adjacent in cancer genomes [13, 26, 5, 22], in sequenced tumor samples. In the more recent study[1] a novel methodological framework `RCK` was proposed for reconstruction of cancer genomes *karyotype graphs*, taking into account both the CNA and NA signals, the non-haploid nature of both the reference and the derived genomes, and, when applicable, possible sample heterogeneity. The karyotype graph provides a much more accurate and complete description of the rearranged cancer genome than either of CNA or NA profiles on their own, but it falls short of describing the actual linear/circular structure of the rearranged chromosomes, and instead provides an alignment of the rearranged chromosomes onto the reference.

Here we consider the problem of inferring sequential structure of rearranged linear/circular chromosomes in a cancer genome given its karyotype graph representation. In previous studies either general observations about necessary and sufficient conditions on the karyotype graphs structure for the existence of the underlying cancer genome [17, 1], or attempts at extracting information about specific local rearranged structures (e.g., amplicons, complex rearrangements, etc) [3, 16, 4] were made. In the present study we formulate a Eulerian Decomposition Problem (EDP) for a cancer karyotype graph with the objective of finding a covering collection of linear and/or circular paths/cycles (i.e., chromosomes). We derive and prove computational complexities for both general, *min*, and *max* versions of the EDP. We further observe that for a cancer karyotype graph there are often multiple possible Eulerian decompositions which presents ambiguities for chromosomal sequential structures inference. We note that similar non-uniqueness of traversals that constitute genome graph decompositions has also been previously observed in the area of genome assembly [11]. We then formulate a Consistent (i.e., shared across all possible minimal Eulerian decompositions) Contig Covering Problem (CCCP) and subsequently present a novel algorithm `CCR`, for <u>C</u>onsistent <u>C</u>ontig <u>R</u>ecovering, capable of solving the CCCP in polynomial time.

We then apply `CCR` method to the karyotype graphs of 17 primary and metastatic prostate cancer samples. We find that `CCR` effectively finds long, unambiguous paths shared across all possible graph decompositions thus reliably inferring cancer contigs. We observe that for these cancer contig inferences, the N50 sizes of the obtained results reaches chromosomal scale, and the total number of contigs is within an order of magnitude of the number of chromosomes in the underlying genomes, whereas a naive algorithm fragments the reconstruction into thousands of contigs and appreciably smaller contig N50 sizes. We thus demonstrate that `CCR` effectively advances the frontier for the analysis of rearranged cancer genomes, allowing for a more detailed and comprehensive studies of structurally altered cancer chromosomes. A proof-of-concept implementation of `CCR` is available at github.com/izban/contig-covering and will be later integrated into the `RCK` software package available at github.com/aganezov/RCK.

## Methods

### Rearranged cancer genomes and interval adjacency graphs

A *segment s* = [*s*^*t*^, *s*^*h*^] is a continuous part of the reference chromosome with *extremities s*^*t*^ and *s*^*h*^ determining its beginning and end respectively. We assume that every segment appears exactly once in the reference, and segments do not overlap. An *adjacency a* = {*p, q*} connects two segments’ extremities *p* and *q* determining a transition between adjacent segments along the (derived) chromosome. We assume that a derived cancer genome Q comes from the reference genome R via large scale rearrangements and can contain both linear and circular chromosomes. For a set *S*(R) of segments in the reference R and a set *S*(Q) of segments in genome Q we naturally have *S*(Q) ⊆ *S*(R) (i.e., “building blocks” that the derived genome is comprised of originate from the reference).

An *Interval Adjacency Graph* (IAG), or *karyotype graph, G*(*S, A*)= (*V, E*) is a graph built on a set *S* of segments and a set *A* of adjacencies. A set *V* = {*s*^*t*^, *s*^*h*^ | *s* ∈ *S*} of vertices represents extremities of segments in a set *S*. A set *E* of edges is comprised of two types of undirected edges: *segment edges E*_*S*_ = {{*s*^*t*^, *s*^*h*^} | *s* ∈ *S*}, encoding segments, and *adjacency edges E*_*A*_ = {{*p, q*} | {*p, q*}∈ *A*}, encoding adjacencies between segments extremities (**Figure 1a**).

**Figure 1:**
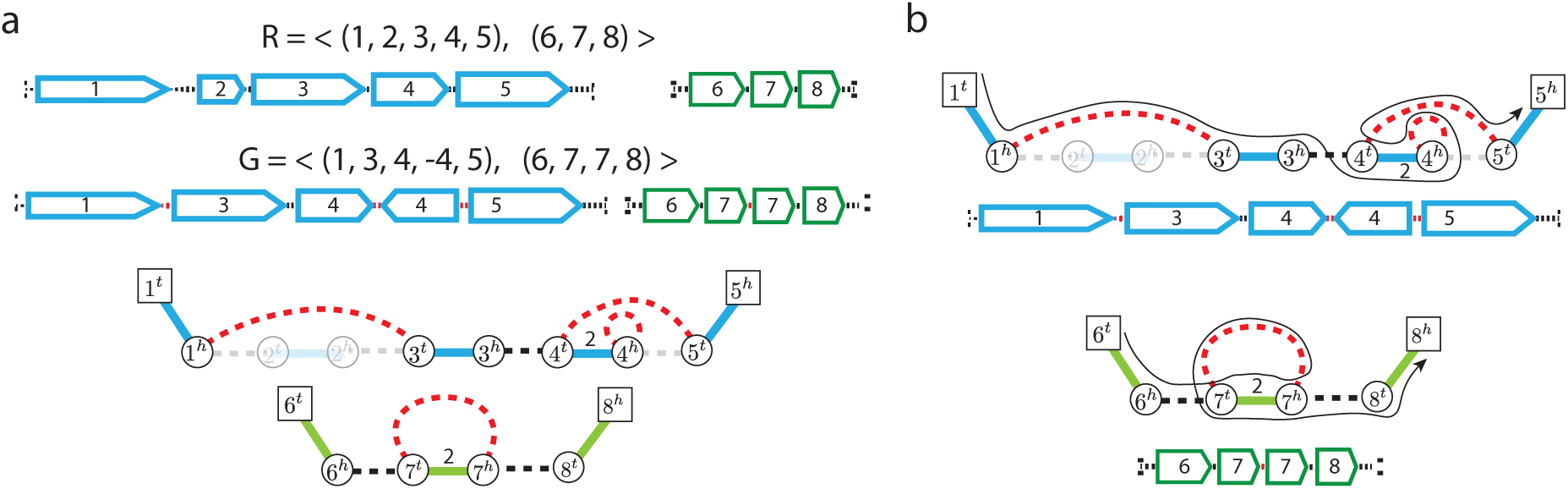
Rearranged genomes and Interval Adjacency Graphs. **a)** Reference and derived genomes R and Q both depicted as sequences of oriented segments and a weighted Interval Adjacency Graph determined by a derived genome Q. In the IAG segment edges are shown as solid, with colors corresponding to distinct chromosomes, and adjacency edges are shown as dashed, with reference adjacencies shown in black, and novel adjacencies depicted in red. Edges with copy number 0 are shown as faded, copy numbers of 1 are omitted for clarity. **b)** Rearranged linear chromosomes in the mutated genome Q depicted as both sequences of oriented segments and as segment/adjacency edge-alternating paths in the IAG’s connected components.

A linear chromosome corresponds to a segment/adjacency edge-alternating path in the IAG that starts with a segment edge and ends with a segment edge (**Figure 1b**). A circular chromosome correspond to a segment/adjacency edge-alternating cycle in the IAG. The number of times every segment/adjacency edge is traversed (in either direction) across all chromosomes in the genome Q corresponds to the segment/adjacency *copy number* (i.e., the number of times segment/adjacency is present in the observed genome).

For an IAG *G* = (*V, E*) we define a multiplicity function *μ* : *E* → ℕ_≥0_ encoding respective segment/adjacency copy number, and we call respective IAG *G* = (*V, E, μ*) *weighted*. The weighted IAG *G* is *positive* if *μ*(*e*) ≥ 1 for all *e* ∈ *E*. Further, since we can trivially derive a positive IAG *G* by removing all edges with with multiplicity 0, we assume that IAG *G* is positive unless explicitly specified otherwise.

For a vertex *v* ∈ *V* we define *e*_*S*_ (*v*) ∈ *E*_*S*_ as a segment edge incident to *v* and *E*_*A*_(*v*) ⊆ *E*_*A*_ as adjacency edges incident to *v*. We also define for all *e* ∈ *E*_*A*_ an auxiliary function *l*(*e*) that outputs 2 if an adjacency edge *e* is a loop, and 1 otherwise.

We define *copy number excess x*(*v*) on a vertex *v* ∈ *V* as follows:

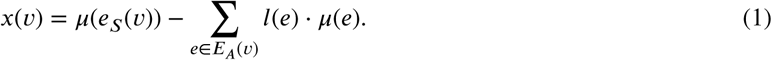

Given that both segment and adjacency edges can have positive multiplicities, for any set *E*′ of edges we define *μ*(*E*′)= ∑_*e*∈*E*′_ *μ*(*e*).

### Interval Adjacency Graph decompositions

For an IAG *G* = (*V, E, μ*) we define *Eulerian decomposition* (ED) as collection *D* = *P* ∪ *C* of segment/adjacency edge-alternating paths *P* and cycles *C* (i.e., linear and circular chromosomes in cancer genome) such that every segment/adjacency edge *e* is present in *D* exactly *μ*(*e*) times. Below we outline the known necessary and sufficient condition for an Eulerian decomposition to exist in an IAG:

#### Lemma 1.

IAG *G* = (*V, E, μ*) has an Eulerian decomposition if and only if for all *v* ∈ *V* holds *x*(*v*) ≥ 0. [20, 17, 1]

We call IAG *G decomposable* if there exist an Eulerian decomposition *D* of *G* and assume that the considered IAG is decomposable unless explicitly stated otherwise.

We define by *C*_+_(*G*) a set of connected components in *G* such that for every *c* ∈ *C*_+_(*G*) there exist a vertex *v* ∈ *c* : *x*(*v*) > 0. We define by *C*_0_(*G*) a set of connected components in *G* such that for every *c* ∈ *C*_0_(*G*) for every vertex *v* ∈ *c* : *x*(*v*)= 0.

As the number of chromosomes can play a crucial role in cancer development and progression we now observe the previously proven result on how the karyotype graph determines the quantity of chromosomes in the depicted cancer genome:

#### Lemma 2.

*For an IAG G* = (*V, E, μ*) *the number* |*P*| *of paths P in any Eulerian decomposition D of G is determined as follows* [1]:

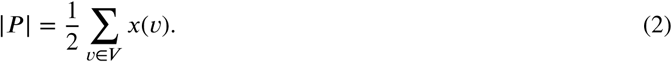

Below we observe the problem of finding the Eulerian decomposition of an IAG which in turn determines the collection of linear/circular rearranged cancer chromosomes:

#### Problem 1

(IAG Eulerian Decomposition Problem, **EDP**). *Given an IAG G find some Eulerian decomposition D of G.*

#### Theorem 1.

***EDP*** *can be solved in O* (*μ*(*E*)) *time.*

*Proof.* We solve the problem for each connected component *c* ∈ *G* separately.

At first, we show how to find an ED of the component *c* from *C*_0_(*G*). In this case, *x*(*v*) = 0 for all *v* ∈ *c*, so the numbers of adjacent segment and adjacency edges (with their multiplicities) are equal in each vertex. Thus, in each vertex we can match each segment edge with some adjacency edge. Since each edge has two matched edges of opposite type, one in each endpoint, these matches represent a set of alternating cycles, which is exactly the desired Eulerian decomposition.

Now, consider a component *c* from *C*_+_(*G*). We modify graph *G* so that *c* becomes the component *c*′ ∈ *C*_0_(*G*′): for each vertex *v* with *x*(*v*) > 0 we add supplemental adjacency loops *l* = {*v, v*} with 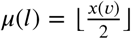; and for the vertices *v*_1_, *v*_2_, …, *v*_2*k*_ which are left with *x*(*v*) = 1 we add supplemental adjacency edges *e*_*i*_ = {*v*_2*i*−1_, *v*_2*i*_} with *μ*(*e*_*i*_) = 1. Indeed, there are an even number of vertices with excess 1 since the total copy number excess ∑_*v∈V*_ *x*(*v*) is even, and thus there may exist only an even number of vertices with an odd copy number excess. In this new graph *G*′ we have *x*(*v*) = 0 for any *v* and, thus, we can build ED *D*′ using the algorithm from the previous paragraph. To get ED of *G* we just remove all supplemental edges from *D*′. We removed only adjacency edges, thus each cycle either remains the same or is split into several edge-alternating paths, which start and end with a segment edge. This satisfies the definition of Eulerian decomposition. □

With the proposed efficient algorithm for obtaining and Eulerian decomposition of an IAG and the respective collection of linear/circular cancer chromosomes, we now observe how many of said chromosomes can be present in the obtained decomposition (**Figure 2**).

**Figure 2:**
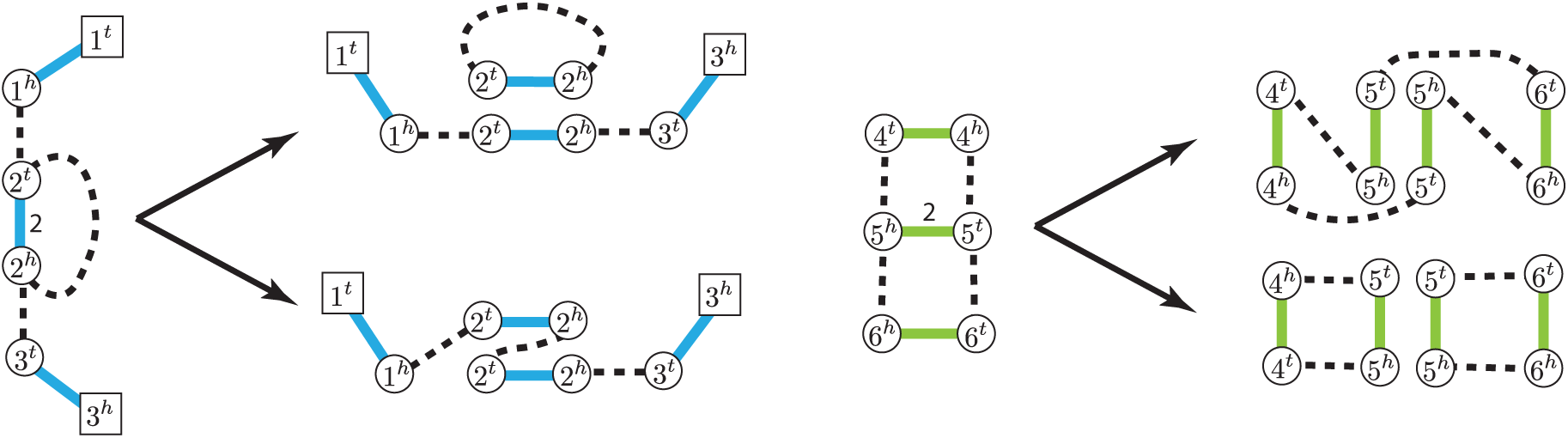
Eulerian Decompositions of Interval Adjacency Graphs. Distinct Eulerian decompositions of different cardinalities of IAG connected components where vertices have a non-zero copy number excess (left) and with all vertices being copy number balanced (right).

Rare circular chromosomes, or *double minutes*, have been previously observed in some cancers [2, 6, 24] and we first consider a case where the number of circular chromosomes in cancer genome is minimal. Below we show the previously known result that for IAG *G* every connected component *c* ∈ *C*_+_(*G*) can be decomposed into paths-only, and every connected components *c* ∈ *C*_0_(*G*) can be covered by a single cycle:

#### Lemma 3.

*For a decomposable IAG G* = (*V, E, μ*) *the size* |*D*| = |*P*| + |*C*| *of any of its minimal (cardinality-wise) Eulerian decompositions is equal to* [1]:

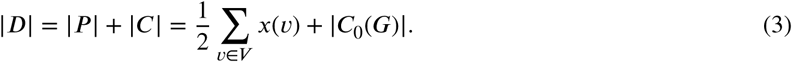

We now described the problem of finding a minimal Eulerian decomposition of an IAG:

#### Problem 2

(IAG Minimal Eulerian Decomposition Problem, **min-EDP**). *Given an IAG G find a* minimal *Eulerian decomposition D of G (i.e., for any Eulerian decomposition D*′ *of G*, |*D*| ≤ |*D*′| ).

#### Theorem 2.

***min-EDP*** *can be solved in O*(*μ*(*E*)^2^) *time.*

*Proof.* At first, using the algorithm from Theorem 1 we obtain some ED *D* of graph *G*. Then, we iteratively merge cycles with paths or cycles with cycles via shared segment edges, until we are left with one cycle in *c* from *C*_0_(*G*), and until there are no cycles in *c* from *C*_+_(*G*). To analyze the overall complexity of the aforementioned workflow for solving the **min-EDP** we first observe that, according to Theorem 1 we can find some ED of *G* in *O*(*μ*(*E*)) time. We then iteratively identify pairs *C, Q* of a cycle *C* and a path/cycle *Q* that share some segment edge *s* and merge them into a new path/cycle *Q*′. At each step a pair *C, Q* can be identified and merged in *O*(*μ*(*E*)), and there can be at most *O*(*μ*(*E*)) such steps, thus bringing the total runtime complexity of the proposed workflow to *O*(*μ*(*E*)^2^).

While the **min-EDP** describes the most parsimonious and biologically relevant scenario, we also observe the opposite case and pose a problem of finding the cancer genome described by a given IAG with the maximum number of circular chromosomes and subsequently prove that it is computationally harder than the **min-EDP**:

#### Problem 3

(IAG Maximal Eulerian Decomposition Problem, **max-EDP**). *Given an IAG G find a* maximal *Eulerian decomposition D of G (i.e., for any Eulerian decomposition D*′ *of G*, |*D*| ≥ |*D*′ |).

#### Theorem 3.

***max-EDP*** *is NP-hard.*

*Proof.* The problem of checking if a simple undirected graph *G* = (*V, E*) has a *K*_3_ edge partition (i.e., a set *E* of edges can be partitioned into non-intersecting subsets *E*_1_, *E*_2_, …, *E*_*n*_ such that each *E*_*i*_ generates a subgraph *G*_*i*_ of *G*, where *G*_*i*_ is isomorphic to a complete graph *K*_3_) is NP-complete [9]. Let us show that **max-EDP** is NP-hard by reducing the problem of *K*_3_ edge partition to **max-EDP**.

Suppose that in the *K*_3_ edge partition problem we are given an graph *G* = (*V, E*), with deg(*v*) as a degree of a vertex *v* ∈ *V*. We note that ∀*v* ∈ *V* we have deg(*v*) ≡ 0 (mod 2) as otherwise *K*_3_ edge partitioning of *G* does not exist. Now we build a new graph *G*′ = (*V* ′, *E*′) from *G* for the **max-EDP** instance as follows: (i) add two vertices *v*^*h*^, *v*^*t*^ ∈ *V* ′ for each *v* ∈ *V* ; (ii) add an adjacency edge *e* = {*v*^*h*^, *u*^*h*^} with *μ*(*e*) = 1 for each edge {*u, v*} ∈ *E*; (iii) add segment edges *s* = {*v*^*h*^, *v*^*t*^} with *μ*(*s*) = deg(*v*); (iv) add adjacency loops *l* = {*v*^*t*^, *v*^*t*^} with *μ*(*l*) = deg(*v*)/2. The example of such transformation is shown in **Figure 3**.

**Figure 3:**
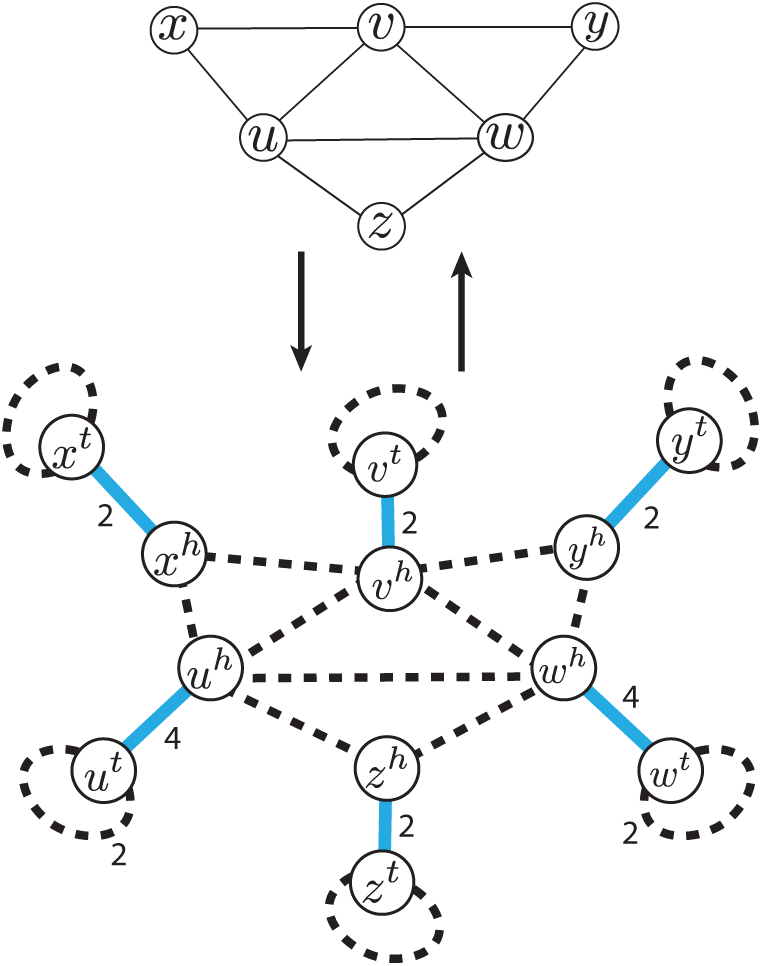
Reduction of *K*_3_ covering problem to the instance of **max-EDP**. Transformation of the regular undirected graph *G* into IAG *G*′ and back with *K*_3_ decomposition in *G* corresponding to segment-adjacency edge-alternating cycles in *G*′.

There exists a bijection between cycles in *G* and segment/adjacency edge-alternating cycles in *G*′. Since the smallest possible cycle in *G*′ corresponds to *K*_3_ in *G*, the solution in **max-EDP** corresponds to finding exactly the partition of *G* into *K*_3_, if it exists. Because the described reduction can be computed in linear time, the **max-EDP** is NP-hard, as is the *K*_3_ edge partition problem. □

### Consistent Contig Covering Problem

We can assume that minimal Eulerian decomposition(s) are the most likely ones, as they describe the simplest possible (composition-wise) representation of a cancer genome. When no additional information is available and there exist several minimal Eulerian decompositions of an observed IAG it is unclear how to select the true one (**Figure 4a**). To address this challenge we consider the inference of sub-paths/cycles that are present across all possible minimal Eulerian decompositions of a considered IAG.

**Figure 4:**
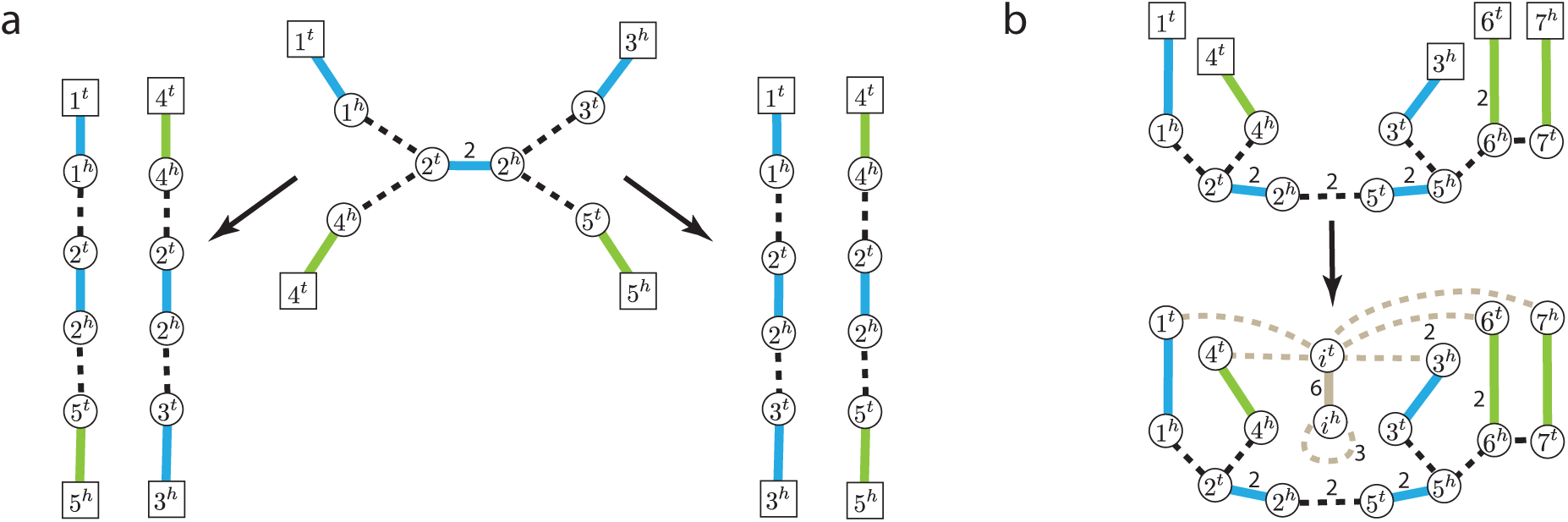
Eulerian decompositions and contigs covering inference for Interval Adjacency Graphs. **a)** Two distinct contig coverings corresponding to minimal Eulerian decompositions of the same IAG. **b)** Transformation of IAG *G* with vertices with copy number excess (i.e., telomere vertices) into a an IAG *G*′ with no telomere vertices. Added supplemental segment and adjacency edges are shown in grey.

For IAG *G* = (*V, E, μ*) a collection *T* = *P* ∪ *C* of segment/adjacency edge-alternating paths/cycles is called a *contig covering* of *G* if (i) every segment edge *e* ∈ *E*_*S*_ is present exactly *μ*(*e*) times across all elements of *T* (i.e., *μ*(*e*) = *μ*_*T*_ (*e*)), (ii) for every adjacency edge *e* ∈ *E*_*A*_ we have *μ*(*e*) ≥ *μ*_*T*_(*e*), where *μ*_*T*_ is the copy number function on edges determined by *T*.

A contig covering of a given decomposable IAG represents (sub)parts of the full linear/circular chromosomes in Eulerian decomposition(s) and we note that an Eulerian decomposition is a special case of a contig covering.

As simple collection of individual segments with their multiplicities being taken into account represents a contig covering which we call *primitive*. Below we consider a problem of finding contig coverings that contain longer paths/cycles and thus represent more contiguous parts of rearranged cancer chromosomes. For IAG *G* = (*V, E, μ*) and its contig covering *T* we define *contiguity discordance* ‖*G* − *T* ‖ as follows:

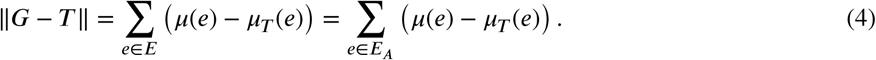

We note, that for IAG *G* = (*V, E, μ*) and any Eulerian decomposition *D* of *G* we naturally have ‖*G* −*D*‖ =0 (i.e., all adjacencies, including multiplicities, are present in *D*) and for a primitive decomposition 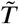 we have 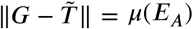.

Since one IAG can have multiple minimal Eulerian decompositions we now formulate a problem of finding the “longest” consistent contigs shared across all minimal Eulerian decompositions (i.e., longest uniquely determined contigs across all possible linear/circular/mixed genomes that the given IAG determines) as follows:

#### Problem 4

(IAG Consistent Contig Covering Problem, **CCCP**). *Given IAG G* = (*V, E, μ*) *find a contig covering T* = *P* ∪ *C of G such that for every minimal Eulerian decomposition D of G every path p* ∈ *P and cycle c* ∈ *C is present in D, and the contiguity discordance* ‖*G* − *T*‖ *is minimized.*

#### Theorem 4.

***CCCP*** *can be solved in O*(*μ*(*E*)^2^) *time.*

*Proof.* At first, we assume that the graph is connected. Otherwise, we consider its connected components separately. Below we present the `CCR` algorithm capable of solving **CCCP** in *O*(*μ*(*E*)^2^). An overview of the `CCR` workflow is shown in **Figure 5**.

**Figure 5:**
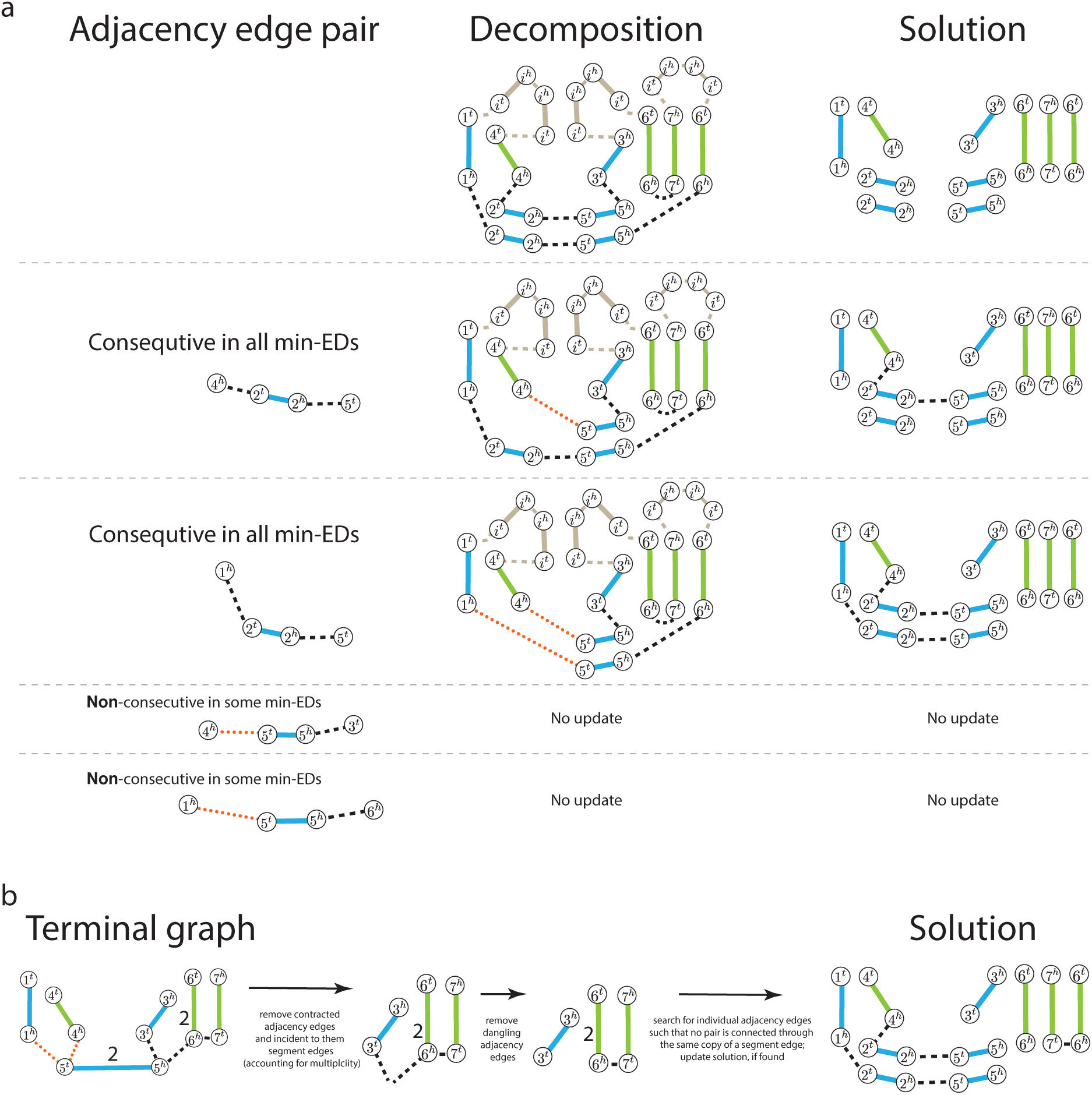
`CCR` workflow on IAG presented in Figure 4b. **a** Iterative Step 3 in `CCR` workflow: processing of pairs of consecutive adjacency edges in the min-ED, contraction of pairs that are present in all min-EDs, updates of the solution. Supplemental (both segment and adjacency) edges are shown in grey, contracted adjacency edges are shown as dotted orange. **b)** Processing of the terminal graph obtained after iterative Step 3 execution with removal of contracted adjacency edges, incident to them segment edges, and dangling edges after that. Search for a collection *L* of adjacency edges in which no pair {*e, f*} share the same copy of a segment.

We start constructing the solution *T* by setting it to the primitive contig covering 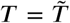 and then iteratively add adjacency edges to it as described below:

1. Transform the initial graph *G* = (*V, E, μ*) into a graph *G*′ which minimal Eulerian decomposition is a single cycle, i.e., *x*(*v*) = 0 for all vertices *v* ∈ *V*. For that we add the following elements into *G*: a *supplemental* segment edge *i* = {*i*^*t*^, *i*^*h*^}, a *supplemental* adjacency edge {*v, i*^*t*^} for every *v* ∈ *V*, and an adjacency edge-loop {*i*^*h*^, *i*^*h*^} (**Figure 4b**). For every added supplemental adjacency edge *e* = {*v, i*^*t*^} we set the multiplicity *μ*(*e*) = *x*(*v*), for supplemental segment edge *i* we set multiplicity *μ*(*i*) = ∑_*v*_ *x*(*v*), and for supplemental self-loop adjacency edge {*i*^*h*^, *i*^*h*^} we set its multiplicity *μ*(*i*)/_2_.
2. Find some minimal Eulerian Decomposition *D*′ of *G*′ using the algorithm from Theorem 2;
3. Since every (sub)path/cycle present in desired *T* must also be present in the obtained *D*′ (when considering only non-supplemental edges) we consider pairs *b* = (*e, f*) of non-supplemental consecutive adjacency edges in any path/cycle in *D*′. In the beginning every such pair *b* is marked as *unprocessed*. For every unprocessed pair *b* = (*e, f*) we check if there exist another minimal Eulerian decomposition of *G*′ where *e* and *f* are not consecutive. For space considerations we describe in detail how this check is performed in the **Supplementary Methods**. If this pair *b* is not consecutive in some other minimal Eulerian decomposition of *G*′ we mark *b* as *processed* and never return to it again. Otherwise, it belongs to all minimal Eulerian decompositions of *G*′. Suppose that *e* = {*x*^*h*^, *y*^*t*^}, *f* = {*y*^*h*^, *z*^*t*^} and the segment edge *y* = {*y*^*t*^, *y*^*h*^} between *e* and *f* in *D*′. We remove *e, y*, and *f* from *G*′ and insert a new non-supplemental contracted adjacency edge *j* = {*x*^*t*^, *z*^*h*^} between endpoints *x*^*t*^ and *z*^*h*^. We update *D*′ in the same manner: we replace *e, y* and *f* with *j*. We further update *T* by adding non-contracted adjacency edges from {*e, f*} by choosing copies of segments *x, y*, and *z* that do not have adjacency edges incident to them (at vertices *x*^*h*^, *y*^*t*^, *y*^*h*^, and *z*^*t*^), and connecting them via *e* and *f*. We update *T* with only non-contracted adjacency edges because all adjacency information from contracted adjacency edges has already been added to *T*. If there are still unprocessed pairs of adjacency edges in *G*′ we reiterate Step 3.
4. We end up with the *terminal graph G*′ in which no pair of adjacency edges belong to the same path/cycle in any consistent contig covering. Now we want to select the largest (cardinality-wise) collection *L* of non-contracted adjacency edges from *G*′ such that when updating *T* with edges from *L* the collection *T* remains consistent contig covering. We find such collection *L* by solving the maximum-matching problem in the auxiliary graph, see **Supplementary Methods** and **Supplementary Figure S1** for details. After updating *T* with adjacency edges from *L* the collection *T* represents a solution to the **CCCP** instance. We note that the solution found to **CCCP** may not be unique.

Now we analyze complexity of the proposed `CCR` algorithm. In Step 2 min-ED can be found in *O*(*μ*(*E*)^2^) as per Theorem 2. There can be at most *O*(*μ*(*E*)) iterations of Step 3 of the algorithm: in each iteration we either mark some pair of edges as processed, or decrease the number of adjacency edges and increase the number of unprocessed pairs by 2. The check adjacency edge pair in Step 3 is performed in *O*(*μ*(*E*)). Complexity of the selection of non-contracted adjacency edges in the terminal graph is dominated by the Maximum-Matching search, which can be performed in *O*(*μ*(*E*)^2^). Thus, the total complexity of `CCR` algorithm is *O*(*μ*(*E*)^2^).

## Results

### Evaluation and applications

To evaluate the performance of `CCR` we applied it on cancer karyotype graphs originally produced with `RCK` on 17 prostate cancer samples [1]. We note that the karyotypes inferred in the previous study are both clone- and haplotype-specific, but for the purposes of this study we do not consider intra-sample relations between distinct clones that comprise it. We also underscore that the `CCR` algorithm and it’s theoretical framework does not discriminate between haplotype-specific and haploid karyotype graphs, as both are Interval Adjacency Graphs at their core.

Out of the 17 considered cancer samples, 10 were heterogeneous with a 2-clone compositions, and 7 were homogeneous with a single cancer clone present in each one of them. In the original study the clone- and haplotype-specific graphs were inferred with `RCK` which was ran with sample-specific novel adjacencies obtained from the earlier study [7] and with both `Battenberg` [7] and `HATCHet` [27] copy number aberrations (CNA) profile inputs, producing a separate graph for each CNA input. Both `Battenberg` and `HATCHet` analyze read-depth ratios and B-allele frequencies of bulk-sequenced short reads that are aligned and grouped across large genomic fragments. Both methods take into account possible sample heterogeneity with `HATCHet` being capable of analyzing jointly multiple samples coming from the same patient. Both methods produce clone- and allele-specific CNA profiles, that `RCK` takes as part of the input along with novel adjacencies and infers the karyotype graph for underlying rearranged cancer genomes. Thus we run `CCR` on 54 karyotype graphs originally inferred for 27 cancer clones in 17 samples.

Information about the lengths (i.e., sum of lengths of segments with their multiplicities taken into account) of the cancer genomes represented as karyotype graphs, the number of chromosomes in their minimal Eulerian decompositions, and whether or not the considered genomes had a Whole Genome Duplication event in the somatic evolutionary history (as according to the original study [27]) is provided in **Supplementary Table S1**.

We observe that `CCR` was capable of recovering long contigs in the obtained consistent contig covering on all observed cancer karyotype graphs, with the total number of unambiguous contigs being within an order of magnitude from the overall number of chromosomes (**Figure 6a,b**). We also computed the N50 size for the obtained contigs, and observe that on average `CCR` obtains contigs with N50 sizes of 50Mbp or greater, which indicates that the `CCR` is capable of recovering chromosomal scale contigs (**Figure 6c,d**). We note that the contig inference results obtained from karyotype graphs produced with either `Battenberg` or `HATCHet` CNA input for `RCK` were very similar, with the inference on `RCK` +`HATCHet` graphs producing fewer and slightly longer contigs on most samples.

**Figure 6:**
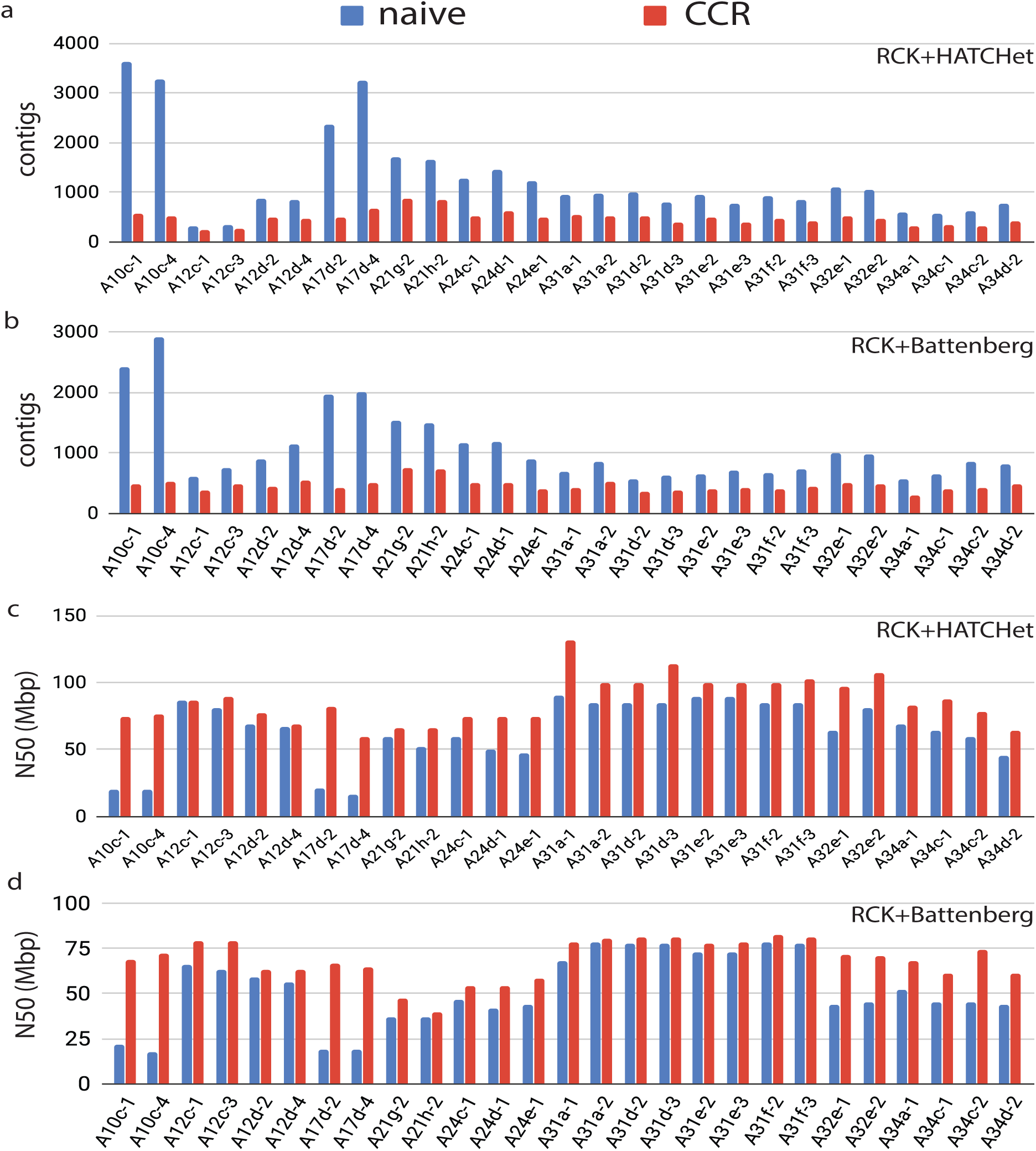
Contigs obtained with both `CCR` (red) and the baseline “naive” contig inference approach (blue) and on haplotype-specific cancer karyotype graphs. **a)** Number of contigs recovered with `CCR` and the “naive” approach from input karyotype graphs obtained with `RCK` on `HATCHet` CNA input. **b)** Number of contigs recovered with `CCR` and the “naive” approach from input karyotype graphs obtained with `RCK` on `Battenberg` CNA input. **c)** N50 metric for contigs recovered with `CCR` and “naive” approach from input karyotype graphs obtained with `RCK` on `HATCHet` CNA input. **c)** N50 metric for contigs recovered with `CCR` and “naive” approach from input karyotype graphs obtained with `RCK` on `Battenberg` CNA input.

We also investigated on whether and if so to what extent the proposed comprehensive `CCR` contig inference algorithm is more efficient than the “naive” approach of recovering contigs, when a contig covering is produced by only including pairs (*a, b*) of consecutive adjacency edges involving a segment edge *s* when their are no other adjacency edges involving *s*. We note that the naive approach, while trivial to derive, has not been previously implemented or utilized and serves as a baseline for assessing the performance of the proposed `CCR` method. We found that `CCR` substantially outperforms the naive approach in both reducing the number of contigs in the produced contig coverings (**Figure 6a,b**) and in the contiguity of the recovered contigs (**Figure 6c,d**).

## Discussion

Inferring rearranged cancer genomes remains a challenging task, but recent advances in inferring clone- and haplotype-specific cancer karyotype graphs genomes have improved our ability to depict and analyze genetic instability in cancer. Our formal graph-based framework and `CCR` algorithm for inferring the sequential structure of cancer genome’s chromosomes from its karyotype graph represents the next logical step in bringing us closer to recovering complete rearranged cancer chromosomes for further analysis.

While our `CCR` algorithm has demonstrated excellent performance on real cancer karyotype graphs with the average N50 of recovered unambiguous contigs exceeding 50Mbp, there are still several avenues for improving and extending the proposed approach. The joint consideration of information from karyotype graphs of several cancer clones that either comprise the same sample or in general belong to the same patient may improve the quality of the obtained contigs, as distinct clones from the same patient have parts of the somatic evolutionary history in common, and thus may share some of the rearranged chromosomal structure. Furthermore, `CCR` would benefit from incorporating, when available, the long-range molecule-of-origin information coming from 3^rd^-generation long/linked reads, that can help resolve some of the ambiguities encountered during the contig inference process. We also note that, while running-time complexity for the minimal Eulerian Decomposition inference step in the `CCR` algorithm is not dominant, a more efficient algorithm for minimal Eulerian Decomposition may exist which would improve upon the the complexity described in Theorem 2.

Overall we believe that `CCR` addresses many of the major limitations in cancer chromosomal structural analysis and can help advance our understanding of both patient/type-specific as well as the overall genetic instability in cancer.

## Conclusions

In the presented work, we analyzed the problem of finding the sequential chromosomal structure of rearranged cancer genomes from their karyotype graph representation. We formulated several variations of the Eulerian decomposition problem (**EDP**) for a given cancer karyotype graph and provided a polynomial-time solution for the most biological relevant **min-EDP** version of it, and proved that the **max-EDP** version is NP-hard.

We observed that the Eulerian decompositions of a given cancer karyotype graph, while representing the sequential structure of the rearranged chromosomes, may not be unique. To combat resulting ambiguities we formulated a consistent contig covering problem (**CCCP**) of inferring a collection of the longest unambiguous (i.e., present in every decomposition) cancer contigs, and presented a novel algorithm `CCR` capable of solving it in polynomial time.

We then evaluated the performance of `CCR` on 54 cancer karyotype graphs obtained with `RCK` on diverse group of 17 prostate cancer samples. Our results consistently showed that `CCR` is capable of decomposing the input karyotype graphs into relatively small collections of long, unambiguous contigs with N50 exceeding 50Mbp in most cases and the total number of contigs within an order of magnitude from the overall number of chromosomes in the considered cancer genomes.

## List of abbreviations

*CNA*: Copy Number Aberrations;
*NA*: Novel Adjacency;
*ED*: Eulerian Decomposition
*EDP*: Eulerian Decomposition Problem;
*CCCP*: Consistent Contig Covering Problem;
*IAG*: Interval Adjacency Graph;
*WGD*: Whole Genome Duplication;

## Availability of data and materials

The datasets generated and/or analysed during the current study are available in the RCK-PUB-DATA repository, https://github.com/aganezov/RCK-pub-data.

## Competing interests

The authors declare that they have no competing interests.

## Funding

SA and MS are supported by the US National Science Foundation (NSF) grant DBI-1350041, US National Institute of Health (NIH) grant R01-HG006677, and Bill and Melinda Gates Foundation. The work of NA is supported by the Government of the Russian Federation through the ITMO Fellowship and Professorship Program. Publication costs are funded via NSF grant DBI-1350041.

## Author’s contributions

SA, NA, and MS conceived of the presented research. SA, NA, and MS formulated described graph-theoretical computational problems. All authors contributed to presented computational results, including the `CCR` method. IZ and VA have implemented the `CCR` algorithm and applied it on prostate cancer karyotype graphs dataset. All authors wrote and approved the manuscript.

## Acknowledgements

Authors would like to think Prof. Benjamin J. Raphael for thoughtful and productive discussions that helped to shape the original `RCK` theoretical framework, the observed graph decomposition research problems and ways to address them. This paper was in part inspired by programming solutions to the problem “Cancer and Chromosome Rearrangements” on the Bioinformatics Contest 2019 https://bioinf.me/en/contest/2019.

