## Supplementary material for "Recovering rearranged cancer chromosomes from karyotype graphs"

### Recovering rearranged cancer chromosomes from karyotype graphs
#### Supplementary material

Sergey Aganezov[1], Ilya Zban[2], Vitaly Aksenov[2,3], Nikita Alexeev[2], and Michael C. Schatz[1,4]

[1]Computer Science Department, Johns Hopkins University, Baltimore, MD, USA
[2]International Laboratory "Computer technologies", ITMO University, Saint Petersburg, Russia
[3]Institute of Science and Technology, Klosterneuburg, Austria
[4]Cold Spring Harbor Laboratory, Cold Spring Harbor, NY, USA

#### Supplemental Methods

##### Adjacency edge pair check in Step 3 in `CCR`

Below, we explain how to perform Step 3 of the `CCR` algorithm: a check that the adjacency edge $e = \{x^h, y^t\}$, the segment edge $y = \{y^t, y^h\}$, and the adjacency edge $f = \{y^h, z^t\}$ are consecutive in all minimal (one-cycle) Eulerian decompositions of $G'$. Below we assume that any edge $e$ with multiplicity $\mu(e)$ can be treated as a collection of $\mu(e)$ parallel edges (i.e., multi-egde).

The following two statements are equivalent: (i) there exists a minimal Eulerian decomposition of $G'$ in which $e$, $y$, and $f$ are non-consecutive; (ii) there exist exactly $\mu(e)$ edges $k = \{y^h, v^t\}$ where $v^t \neq z^t$ such that after the replacement of all triples $e$, $y$, and $k$ with adjacency edge $\{x^h, z^t\}$ any minimal Eulerian decomposition of the resulting graph $\tilde{G}$ consists of one cycle.

The second statement follows from the first one, since we can choose exactly $\mu(e)$ edges which follow $e$ and $y$ in the satisfying Eulerian decomposition of $G$. To prove the other direction we can take any minimal Eulerian decomposition of $\tilde{G}$ and undo the replacements (i.e., replace one added adjacency edge back to three original edges) to get the satisfying Eulerian decomposition in $G'$.

Now instead of checking the existence of a minimal Eulerian decomposition of one cycle in $\tilde{G}$ we can simply check that $\tilde{G}$ is connected because the constraint "$x(v) = 0$ for all $v$" holds by construction: $G'$ satisfies this constraint and each replacement does not change $x(v)$ for any vertex $v$. Thus, we check the existence of $\mu(e)$ edges which replacement produces the connected graph.

Let $K$ be the set of all edges $k_i = \{y^h, v_i^t\}$, where $v_i^t \neq z^t$. We account for each edge $e$ $\mu(e)$ times. Consider three cases:

- $|K| < \mu(e)$. By the Dirichlet's Principle each minimal Eulerian Decomposition must contain three consecutive edges $e$, $y$, and $f$.
- $|K| = \mu(e)$. There exist only one way to choose $\mu(e)$ edges. Thus, we simply replace the only triples that are possible and check the connectivity of the resulting graph.
- $|K| > \mu(e)$. Let $G^d$ be the graph $G'$ after we remove all $\mu(e)$ copies of $e$ and all the edges from $K$. Now we have to choose for each edge $k_i$ how to put it back into $G^d$: either as $k_i = \{y^h, v_i^t\}$ or as $\overline{k_i} = \{x^h, v_i^t\}$, and there should be exactly $\mu(e)$ edges of the second type. Note that we do not remove edge $y$ from $G'$, since we only check the connectivity and $\mu(y) > |K| > \mu(e)$, in other words, $y$ will be in a graph after $\mu(e)$ replacements.

  Suppose that graph $G^d$ has $\ell$ components. Let $C(v)$ be the identifier of the component that contains $v$. Then, the set of components is determined as $\{C(x^h), C(y^t) = C(y^h), C(v_1^t), \dots, C(v_{|K|}^t)\}$. We state that it is possible to connect components by placing edges back in $G^d$ if and only if $|K| \geq \ell - 1$.

If $|K| < \ell - 1$, we cannot connect the graph with $\ell$ components using less than $\ell - 1$ edges. Now, we show how to put the edges back if $|K| \geq \ell - 1$. We consider four cases:

- $C(x^h) = C(y^h)$. We can choose any $\mu(e)$ edges: both $k_i$ and $\overline{k_i}$ connects component $C(v_i^t)$ with component $C(x^h) = C(y^h)$.
- For some $i$, $C(v_i^t) = C(x^h)$. We put $k_i$ in $G^d$ and get $C'(y^h) = C'(x^h) = C'(v_i^t)$. After, we can choose any $\mu(e) < |K|$ edges out of the rest: both $k_i$ and $\overline{k_i}$ connect component $C(v_i^t)$ with component $C'(x^h)$.
- For some $i$, $C(v_i^t) = C(y^h)$. We put $\overline{k_i}$ in $G_d$ instead of $k_i$ and get $C'(y^h) = C'(x^h) = C'(v_i^t)$. After, we can choose any $\mu(e) - 1 < |K|$ edges out of the rest: both $k_i$ and $\overline{k_i}$ connect component $C(v_i^t)$ with component $C'(x^h)$.
- Otherwise, $C(x^h) \neq C(v_i^t)$, $C(y^h) \neq C(v_i^t)$ for any $i$, and $C(x^h) \neq C(y^h)$. Thus, the edges from $K$ connect $C(y^h)$ with $\ell - 2$ different components. Since $|K| \geq \ell - 1$, there exists $p$ and $q$ such that $C(v_p^t) = C(v_q^t)$. We put $k_p$ and $\overline{k_q}$ in $G^d$, and get $C'(y^h) = C'(x^h) = C'(v_p^t)$. After, we can choose any $\mu(e) - 1$ edges out of the rest: both $k_i$ and $\overline{k_i}$ connect component $C(v_i^t)$ with component $C'(x^h)$.

##### Terminal graph processing in Step 4 in CCR

After iterative application of Step 3 in CCR we end up with the terminal IAG $G'$ in which no pair of adjacency edges is present in any of consistent contig coverings. With that being noted we observe that individual adjacency edges may still be added to the solution $T$ and improve the $\|G - T\|$ objective. Here we observe how we select such collection $L$ of adjacency edges to be added to $T$.

First, we observe that we can not select any pair $\{e, f\}$ of adjacency edges in $G'$ that are connected through the same copy of some segment edge. Another way of saying it: every segment edge can be incident to at most one adjacency from $G'$ in $T$. Since contracted adjacency edges determine some collection of original adjacency edges that were already added to $T$ during Step 3 (i.e., already adjacent to contracted adjacency edges), we remove all contracted adjacency edges from $G'$ as well as segment edges incident to them (accounting for multiplicity). After this procedure we can end up with some vertices having only adjacency edges incident to them. We call adjacency edges that have do not have a segment edge on at least one of their extremities *dangling*, and remove them from the $G'$.

**Figure S1: Workflow for selection of a maximum collection $L$ of adjacency edges in the terminal graph in CCR.** Terminal graph $G'$ contains a total of 5 adjacency edges with a total multiplicity of 6. In segment-connectivity graph $\widehat{G'}$ vertices are determined by segment edges in $G'$ and edges are determined by adjacency edges in $G'$. Both vertex and edge multiplicities are shows next to them. In expanded segment-connectivity graph $\overline{\widehat{G'}}$ every pair of adjacency vertices $x_1, x_2$ are determined by adjacency edge $x$ in $\widehat{G'}$. For every vertex $u$ with multiplicity $\mu(u)$ in $\widehat{G'}$ $\mu(u)$ segment vertices are added into $\overline{\widehat{G'}}$. Edges in $\overline{\widehat{G'}}$ are determined by either edges in $\widehat{G'}$ (i.e., edges between paired adjacency vertices; e.g., $\{a_1, a_2\} \leftrightarrow a$) or by adjacent vertices in $\widehat{G'}$ (e.g., $\{1, 2\} \leftrightarrow \{a_1, 1\}, \{a_2, 2_1\}, \{a_2, 2_2\}, \{a_2, 2_3\}$). Maximum-matching (edges shown in bold) in $\overline{\widehat{G'}}$ determines a collection $\overline{\widehat{L}} = [\{a_1, a_2\}, \{b_1^2, b_2^2\}, \{d_1, d_2\}]$ of paired adjacency vertices (shown in green) that in turn determine a collection $\widehat{L} = [a, b, d]$ in $\widehat{G'}$, which ultimately determines a collection $L = [\{1^h, 2^t\}, \{2^h, 3^t\}, \{6^h, 2^t\}]$ of adjacency edges in $G'$.

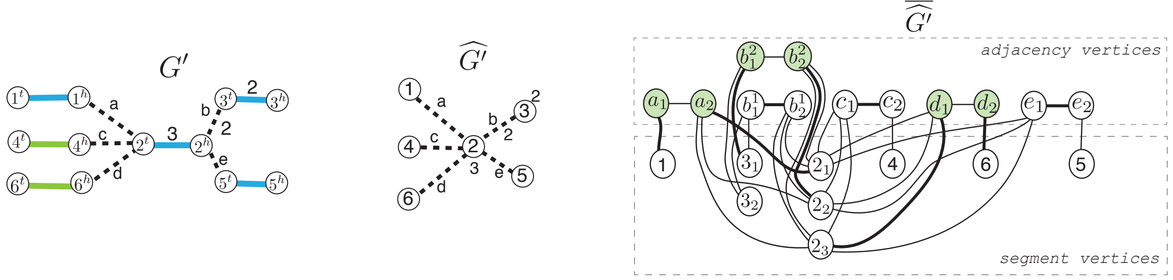

Now in the "cleaned" terminal graph $G'$ we need to select a collection $L$ of adjacency edges still complying with the constraint that no two adjacency edges may be connected through the same copy of some segment edge. To solve this problem we first obtain a segment-connectivity graph $\widehat{G'}$ in which vertices are determined by segment edges in $G'$, and edges are determined by adjacency edges in $G'$. We also retain multiplicity information $\mu(e)$ for every segment/adjacency $e$ and record it for respective vertices ($\mu(v)$) and edges ($\mu(e)$) in $\widehat{G'}$ (Supplementary Figure S1). Original problem of finding $L$ can now be reformulated as a problem of finding the largest (cardinality-wise) collection $\widehat{L}$ of edges in $\widehat{G'}$ such that every vertex $v$ is present at most $\mu(v)$ across all edges in $L$ with edges multiplicities taken into account.

To find such collection $\widehat{L}$ we transform the segment-connectivity graph $\widehat{G'}$ into the expanded segment-connectivity graph $\overline{\widehat{G'}}$. Vertices $V$ in $\overline{\widehat{G'}}$ are comprised of two types of vertices: (i) *segment vertices* $V_S$, where every vertex $v$ in $\widehat{G'}$ corresponds to $\mu(v)$ vertices $v_1, v_2, \dots, v_{\mu(v)}$ vertices in $V_S$; (ii) paired *adjacency vertices* $V_A$, where every edge $a = \{v, u\}$ in $\widehat{G'}$ corresponds to $2\mu(a)$ vertices $a_v^1, a_v^2, \dots, a_v^{\mu(a)}$ and $a_u^1, a_u^2, \dots, a_u^{\mu(a)}$ respectively. Edges $E$ in $\overline{\widehat{G'}}$ are determined as follows: (i) for every edge $a = \{v, u\}$ in $\widehat{G'}$ add edges $\{a_v^i, a_u^i\}$ for all paired adjacency vertices determined by $a$ to $E$; (ii) for every edge $a = \{v, u\}$ in $\widehat{G'}$ add edges $\{a_v^i, v_i\}$ and $\{a_u^j, u_j\}$ for all $i \in \mu(v)$ and $j \in \mu(u)$ to $E$.

We now search for a maximum matching $M$ in the constructed $\overline{\widehat{G'}}$ and then observe edges involved in $M$. If for a paired adjacency vertices $a_u^i, a_v^i$ both of them are present in $M$ in different edges (i.e., connected to segment vertices) we considered such paired adjacency vertices as well as the respective copy $i$ of the edge $a$ in $\widehat{G'}$ *selected*. If for some paired adjacency vertices $a_u^i, a_v^i$ only one of them (e.g., $a_u^i$) is involved in $M$ it is easy to see that we can construct another maximum matching by substituting an edge $\{a_u^i, x\}$ from $M$ with $\{a_u^i, a_v^i\}$. Selected edges in paired adjacency vertices and respective edges in $\widehat{G'}$ determine he searched collection $L'$, which in turn determines the collection $L$. We note that $|M| = |L| + \mu(E)$, where $\mu(E)$ is the combined multiplicity of adjacency edges in $G'$.

We prove that the obtained collection $L$ is the optimal one by contradiction. Let us assume that there exists another collection $\tilde{L}$ of adjacency edges in $G'$ such that (i) no pair of adjacency edges in $\tilde{L}$ shared the same copy of any segment edges; (ii) $|L| < |\tilde{L}|$. We then observe the graph $\overline{\widehat{G'}}$ and select the following edges in it: (i) for every copy $i$ of an adjacency edge $e = \{u, v\}$ from $G'$ not in $\tilde{L}$ we select an edge $\{a_u^i, a_v^i\}$; (ii) for every copy $i$ of an adjacency edge $e = \{u, v\}$ from $\tilde{L}$ we select an edge $\{a_u^i, u_i\}$ and an edge $\{a_v^i, v_i\}$. Selected edges comprise a matching $\tilde{M}'$ in $\overline{\widehat{G'}}$ as every adjacency vertex participated in exactly one selected edge, and no segment vertex was selected more than once with an edge, as otherwise some adjacency edges from $\tilde{L}$ would share a segment edge. We observe that our selection matching $\tilde{M}$ has a cardinality of $\mu(E) + |\tilde{L}|$, where $\mu(E)$ is the combined multiplicity of edges in $G'$, and $|M'| > |M|$ (since $|L| < |\tilde{L}|$), which contradicts the fact that $M$ is a maximum matching.

### Cancer Genome Characteristics

**Table S1:** Statistics for input haplotype-specific cancer genome karyotype graphs obtained with `RCK` with either `HATCHet` or `Battenberg` copy number aberrations (CNA) profiles: presence/absence of Whole Genome Duplication (WGD) events in the somatic evolutionary history (as predicted by CNA input method), the total length (i.e., combined length of the genomic segments with their multiplicities taken into account) of the rearranged genomes represented by the karyotype graphs, and the number of both linear and circular chromosomes in the minimal Eulerian decomposition of the respective karyotype graphs.

| **sample** | WGD | input CNA for `RCK` | **id** | **genome length** | **# of chr** |
|---|---|---|---|---|---|
| A10c | - | `HATCHet` | A10c-1 | 6913472300 | 55 |
| A10c | - | `HATCHet` | A10c-4 | 6477409158 | 49 |
| A10c | - | `Battenberg` | A10c-1 | 6806518228 | 51 |
| A10c | - | `Battenberg` | A10c-4 | 6074369908 | 53 |
| A12c | - | `HATCHet` | A12c-1 | 5830616622 | 50 |
| A12c | - | `HATCHet` | A12c-3 | 6046116477 | 49 |
| A12c | + | `Battenberg` | A12c-1 | 11921775296 | 97 |
| A12c | + | `Battenberg` | A12c-3 | 12195193992 | 100 |
| A12d | - | `HATCHet` | A12d-2 | 5911876634 | 46 |
| A12d | - | `HATCHet` | A12d-4 | 6226722034 | 49 |
| A12d | - | `Battenberg` | A12d-2 | 6086938174 | 47 |
| A12d | - | `Battenberg` | A12d-4 | 6461954615 | 51 |
| A17d | - | `HATCHet` | A17d-2 | 6127215511 | 69 |
| A17d | - | `HATCHet` | A17d-4 | 6153796213 | 70 |
| A17d | - | `Battenberg` | A17d-2 | 5739088237 | 61 |
| A17d | - | `Battenberg` | A17d-4 | 6199585444 | 81 |
| A21g | - | `HATCHet` | A21g-2 | 6673539214 | 56 |
| A21g | - | `Battenberg` | A21g-2 | 6689196116 | 56 |
| A21h | - | `HATCHet` | A21h-2 | 6676278130 | 56 |
| A21h | - | `Battenberg` | A21h-2 | 6670699415 | 60 |
| A24c | - | `HATCHet` | A24c-1 | 6075976625 | 51 |
| A24c | - | `Battenberg` | A24c-1 | 6076959656 | 49 |
| A24d | - | `HATCHet` | A24d-1 | 6068500046 | 49 |
| A24d | - | `Battenberg` | A24d-1 | 5873433192 | 48 |
| A24e | - | `HATCHet` | A24e-1 | 6091325676 | 51 |
| A24e | - | `Battenberg` | A24e-1 | 6006670677 | 48 |
| A31a | + | `HATCHet` | A31a-1 | 11178018498 | 85 |
| A31a | + | `HATCHet` | A31a-2 | 11122817615 | 88 |
| A31a | + | `Battenberg` | A31a-1 | 10370908960 | 81 |
| A31a | + | `Battenberg` | A31a-2 | 11248056960 | 88 |
| A31d | + | `HATCHet` | A31d-2 | 9190788247 | 88 |
| A31d | + | `HATCHet` | A31d-3 | 9758365321 | 77 |
| A31d | + | `Battenberg` | A31d-2 | 10370908960 | 74 |
| A31d | + | `Battenberg` | A31d-3 | 9571306435 | 77 |
| A31e | + | `HATCHet` | A31e-2 | 10936617621 | 87 |
| A31e | + | `HATCHet` | A31e-3 | 9459810642 | 75 |
| A31e | + | `Battenberg` | A31e-2 | 8851146568 | 71 |
| A31e | + | `Battenberg` | A31e-3 | 10172668156 | 80 |
| A31f | + | `HATCHet` | A31f-2 | 10951324664 | 87 |
| A31f | + | `HATCHet` | A31f-3 | 9536558107 | 75 |
| A31f | + | `Battenberg` | A31f-2 | 9804346269 | 78 |
| A31f | + | `Battenberg` | A31f-3 | 10471463652 | 84 |
| A32e | + | `HATCHet` | A32e-1 | 10634418132 | 86 |
| A32e | + | `HATCHet` | A32e-2 | 10077348942 | 77 |
| A32e | + | `Battenberg` | A32e-1 | 10310217848 | 80 |
| A32e | + | `Battenberg` | A32e-2 | 11060253950 | 88 |
| A34a | - | `HATCHet` | A34a-1 | 5882675096 | 45 |
| A34a | - | `Battenberg` | A34a-1 | 5878113411 | 45 |
| A34c | - | `HATCHet` | A34c-1 | 5629706236 | 46 |
| A34c | - | `HATCHet` | A34c-2 | 5956408408 | 46 |
| A34c | - | `Battenberg` | A34c-1 | 5832104111 | 49 |
| A34c | - | `Battenberg` | A34c-2 | 5828229774 | 48 |
| A34d | - | `HATCHet` | A34d-2 | 5747700215 | 45 |
| A34d | - | `Battenberg` | A34d-2 | 5774450082 | 47 |
